# Novel model combining intrinsic and learned behaviours captures divergent effects of dopaminergic drugs on different types of motivation

**DOI:** 10.64898/2026.01.07.698166

**Authors:** Madeleine Bartlett, P. Michael Furlong, Terrence C. Stewart, Jeff Orchard, Megan G Jackson

**Author notes:** Senior author (Jeff Orchard, Megan G Jackson). Technical contact (Madeleine Bartlett).

## Abstract

Motivation-based symptoms occur in an array of neurological and neurodegenerative disorders, including Schizophrenia and Parkinson’s Disease. Unfortunately, there is, as yet, no agreed treatment approach. To establish a better understanding of motivation, and so highlight potential avenues for treatment, motivation is often investigated using operant behavioural paradigms such as the Effort for Reward (EfR) task. Performance on these tasks is thought to be influenced by reinforcement learning (RL) mechanisms, such that disorders of motivation can be described in terms of altered interactions with RL processes. Recently, foraging behaviour has been increasingly adopted as an ethologically valid approach to investigating cognitive mechanisms, including motivation. One example of this is the recently developed Effort Based Foraging task (EBF). Foraging behaviour, unlike the strictly controlled and contrived operant behavioural paradigms, involves a series of complex behavioural processes, each potentially driven by different cognitive processes. It is therefore important to identify which portion of the foraging behavioural sequence is driven by the same RL mechanisms as classical operant behavioural paradigms, and therefore can be used to investigate motivation. In this work we set out to establish whether the same RL mechanisms could be used to account for behaviours observed in both EfR and EBF tasks. We identified where, within the EBF task, RL mechanisms were no long sufficient and developed a novel hybrid model of the behaviour. This model successfully accounted for external influences on motivation, including previously unpredicted effects of clinically used dopaminergic drugs. This work reveals that motivation in complex, naturalistic tasks cannot be fully explained by learning-based models alone. Incorporating intrinsic behavioural drives may be needed as neuroscience moves toward more ethological behavioural assays.

## INTRODUCTION

To address challenges associated with translation of psychiatric symptoms using animal models, there has been a recent shift in the field of neuroscience towards the adoption of more ethologically valid behavioural paradigms. Foraging behaviour in particular shows promise as a tool for investigating key cognitive processes^1^. It requires a series of complex behavioural processes with relevance to a range of neuropsychiatric and neurodegenerative symptoms including decision making, cognitive flexibility and motivation. To account for behaviours observed via these novel paradigms, computational neuroscientists must face the challenge of capturing behaviours and actions that may be driven by more complex interactions between cognitive mechanisms. Motivation-based symptoms are of particular clinical and scientific interest due to their prevalence across neurological and neurodegenerative disease and lack of agreed treatment approach^2^ ^3,4^.

In animal models, motivation is commonly investigated through established operant behavioural paradigms such as the Effort for Reward task (EfR) and Progressive Ratio task (PR)^5,6^. In these paradigms, an animal is food- or water-restricted and trained over a period of weeks to months to make an effortful number of conditioned responses (e.g., nose poke, lever press) to obtain reward. The decisions the animal makes across the session provide a readout of motivational state. These task conditions ensure robust and specific behavioural output is obtained. However, externally modulating motivational state through food/water restriction and relying only on extrinsic reward may fail to capture deficits in intrinsic, self-initiated behaviours that more closely align to motivational disorders. To capture motivational state in the absence of prolonged training periods or food/water restriction, a novel foraging paradigm, the Effort Based Foraging (EBF) task, was developed^7^. In the EBF task, the mouse is placed in an arena with two main components; a safe and enclosed home area, and a foraging area connected via a tube. The mouse can freely choose to traverse the tube and obtain nesting material from a custom designed ‘nesting material box’ which requires effort to forage. The amount of nesting material foraged is a readout of motivational state. Notably, it does not require training beyond habituation and is not driven by food or water restriction. Task output has been shown to be sensitive to pharmacological modulation of the dopaminergic and serotonergic system^7–9^, as well as to phenotypic disease states where motivational deficit is known to occur^10–13^. Behavioural output was also sensitive to increasing levels of effort, and changes in the aversiveness of the foraging environment^7,14^. As such, this task provides a rapid and sensitive readout of motivational state in the absence of physiological restriction and overtraining as required by traditional approaches.

This departure from a more reductionist approach where researchers attempt to isolate the cognitive function of interest through simplistic environments and restriction allows for better generalisation of findings to natural behaviour, but also presents a number of challenges. In particular, from the perspective of computational neuroscience, more contrived paradigms are designed to isolate cognitive mechanisms, allowing researchers to establish the precise nature of those mechanisms and their responses to systematic perturbations. For example, motivated behaviours are often described in terms of reward-based learning mechanisms such as reinforcement learning (RL)^15^. RL models of behaviour can, therefore, be very useful in providing accounts of behaviours relating to motivation, and of the possible mechanisms underlying disorders of motivations. Ethologically valid foraging behaviours, on the other hand, employ multiple cognitive processes, implying the need to develop larger scale computational models, or specify the portion of the observed behaviour that is being driven by the cognitive mechanism of interest. The importance of the distinction between these two task contexts was highlighted during the validation of the EBF. Xeni et al. ^7^ revealed an effect that seems to contradict predictions from the EfR and PR literature. Namely, within the EfR and PR tasks, drugs that act as dopamine agonists (i.e. methylphenidate and amphetamine) lead to an increase in the frequency of effortful behaviours^9^. As predicted, Xeni et al. ^7^ observed an increase in the amount of material being foraged following methylphenidate administration. However, when mice were administered amphetamine, the researchers observed a decrease in the amount of material foraged. Given the difficulty in treating motivational symptoms clinically, this divergence in drug effect may have critical translational relevance. It is important that these differences are similarly detected at the computational level.

The mesolimbic dopaminergic system consists of dense dopaminergic projections connecting the ventral tegmental area to the ventral striatum of the basal ganglia. This system is thought to play a central role in reinforcement or associative learning, where dopamine is thought to carry the reward prediction error signal. This signal is integral to many mathematical models of RL, including the temporal difference (TD) learning rule(s)^16^. If motivated behaviour in the EBF task is governed by the same RL mechanisms that explain behaviour in operant paradigms, then models based on TDRL should account for behaviour across both task contexts. Conversely, failures of these models would suggest that additional, non-learning-based processes contribute to motivated foraging.

Therefore, in this work, we first assess whether TDRL can account for the behaviours observed in both the EBF and EfR tasks. We demonstrate that TDRL alone is an inadequate model of the foraging task, and that a hybrid model, incorporating both TDRL and the assumption of intrinsically driven behaviour, achieves better prediction accuracy. We next incorporate additional biological details into the TDRL model, in the form of changes to tonic and phasic dopaminergic signalling, to assess whether the novel hybrid model could capture the observed divergence in drug effect between the two tasks. We again observed that the hybrid model is unique in capturing the observed effects of the drugs in the EBF. Finally, we utilised our novel hybrid model to generate dose response predictions to inform future experimental work. Overall, this work demonstrates that models which rely solely on learning mechanisms, without including intrinsic behaviours, cannot capture nuanced effects revealed by foraging paradigms. This has important implications for the way that motivated behaviour is modelled as behavioural assays become more complex.

## RESULTS AND DISCUSSION

### A curriculum model requires over-training in order to solve the EBF task

To investigate whether the EBF task could be learned using only TDRL, we developed a custom environment using the MiniGrid environments^17^ as a base. The bespoke environment is depicted in Figure 1. The agent’s (red triangle) task was to learn to navigate from the home location (top left-hand corner) to the forage location (bottom right-hand corner), forage for the key, and bring the key back to the home location. For the curriculum model, the agent was trained to solve the task using a curriculum learning protocol – the task to be learned is broken into several stages and the agent is trained on each stage sequentially. Further details about the environment and the training protocols are provided in the *Methods* section.

**Figure 1.**
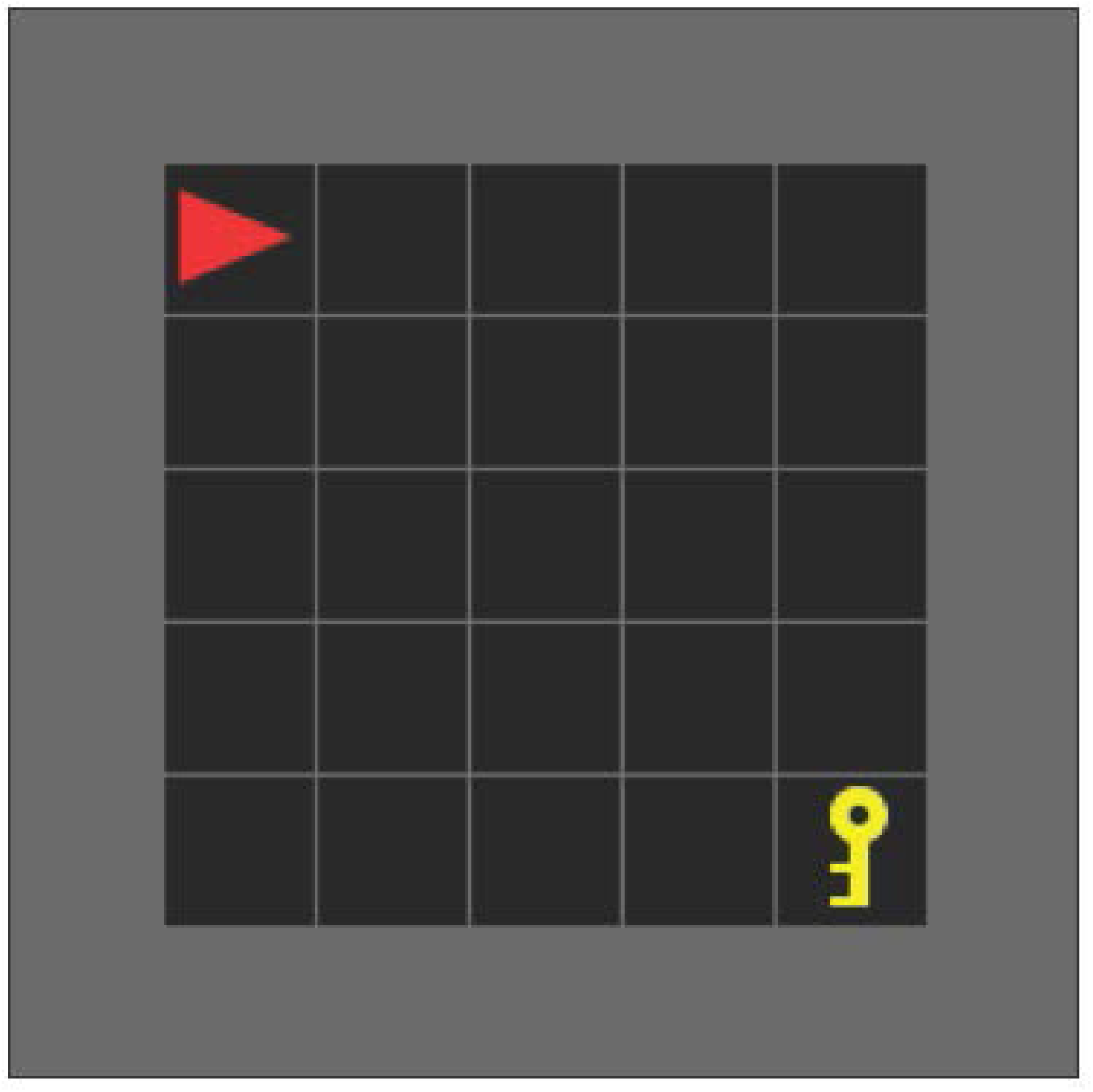
Simulated Effort-Based Foraging environment. Agent is depicted as a red triangle in the top left-hand corner (the home nest location). Forage material is represented as the yellow key object in the bottom right-hand corner. Agent can turn left, turn right, move forward, pick up object or drop object. Environment developed using the Farama Foundation MiniGrid library.

Once a trained agent was obtained, we examined how well it captured the behaviours observed in Xeni et al. ^7^ by testing how the agent responded to changes in the amount of effort required to obtain the reward and in the aversiveness of the environment. Xeni et al. ^7^ found that increasing effort required to forage decreased foraging behaviour. Increasing the aversiveness of the forage area by increasing its size also suppressed foraging behaviour. For increasing effort, we introduced a penalty for the act of foraging (*−*0.5) and multiplied that penalty by *×*1, *×*10, *×*50, and *×*100 to obtain 4 different levels of effort. The environment aversiveness was an analogue of making the environment bigger as in Xeni et al. ^7^ . To simulate this effect, we introduced a penalty of 0, *−*1, *−*5, and *−*10 for being outside of the home location.

The simulated agents follow a similar trend as the real mice in that, as effort increases, the number of rewards obtained decreased (Kruskal Wallis: *H* = 173.474, *df* = 3, *p* =*<* 0.001, *η*^2^ = 0.718, (Effort = 50, *p <* 0.001) (Effort = 100, *p <* 0.001)) (figure 2A). A similar pattern is observed when the environment was made more aversive (Figure 2C) (Kruskal Wallis: *H* = 187.798, *df* = 3, *p* =*<* 0.001, *η*^2^ = 0.779, (Aversion = 5, *p <* 0.001) (Aversion = 10, *p <* 0.001)).

**Figure 2.**
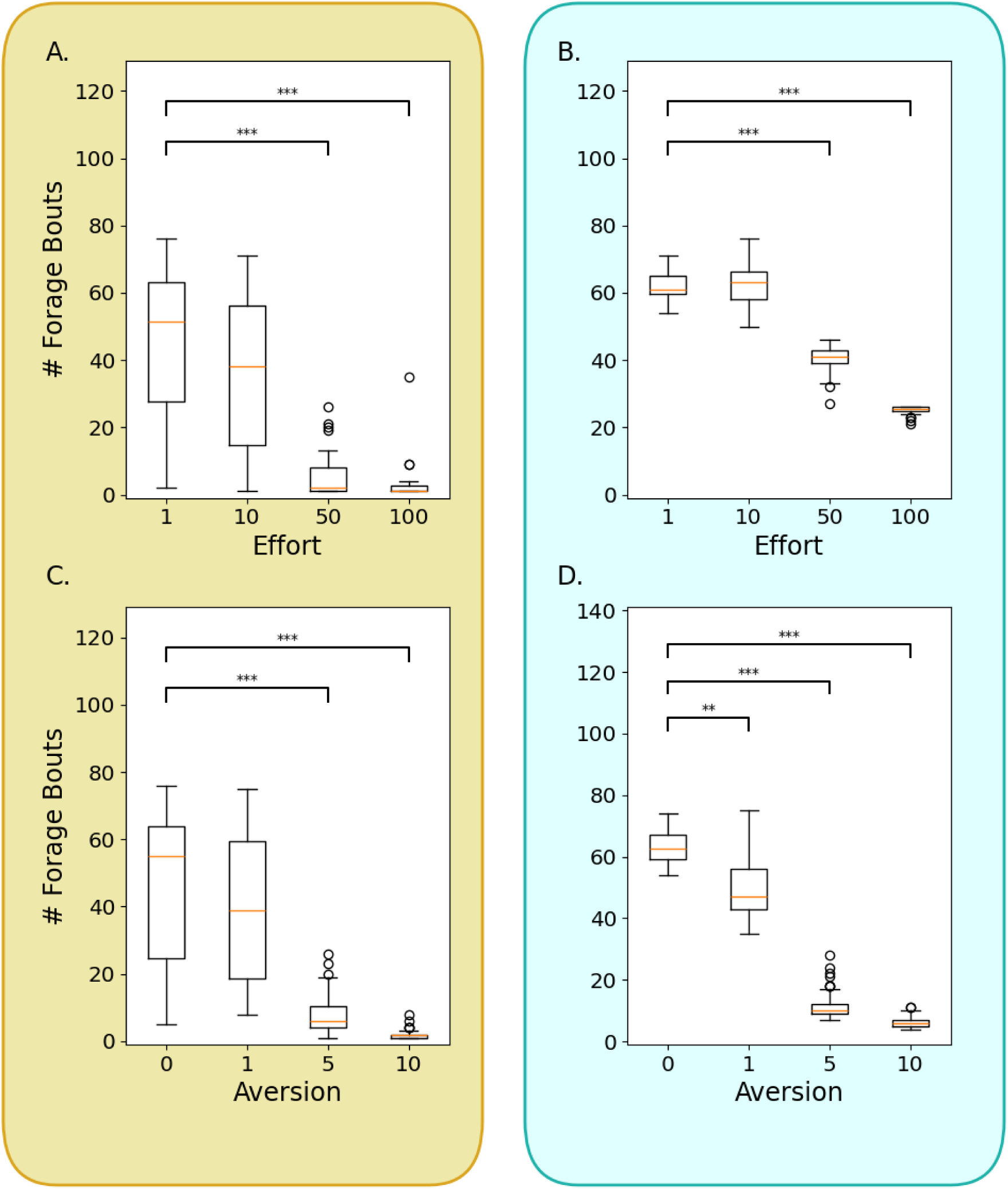
Effort and aversiveness of the environment decreases the number of forage bouts performed by both models. Boxplot showing the number of forage bouts performed by the curriculum model (yellow, left) and the hybrid model (blue, right) over 60 independent trials in each effort and aversion condition. A. & B. Effort, C. & D. Aversion. \**p <* 0.05, \*\**p <* 0.01, \*\*\**p <* 0.001

What we note here is that following the curriculum training protocol did not reliably produce agents that had learned the task. For our purposes, ‘learning the task’ means finding the most efficient path to obtaining bedding and returning it to the home location. We see this behaviour in mice when they make repeated excursions to get bedding material, taking the efficient path once they’ve learned it, and returning the gathered bedding material to a safe/home location. In our experiments, for those agents that did learn the task, in order to observe a substantial change in behaviour, we had to increase the temperature of the softmax used for probabilistically selecting actions from 1 to 10. *Temperature* controls the sharpness of the decision boundary between competing actions such that increasing the temperature, decreases the sharpness. Practically, this has the effect of introducing more opportunities for exploration–selecting actions that are not the learned ‘optimal’ choice. Without this change, the agent’s behaviour was at ceiling and extreme changes to aversion and effort were required to observed any reduction in the number of forage bouts. This illustrates the degree to which the curriculum agent over-learned the task. However, this over-learning was essential to obtaining an agent that successfully performed the foraging task.

One conclusion that can be drawn from this is that the EBF task is likely not entirely solved through reinforcement learning. Rather, part of the task can be learned (i.e. navigating to a forage location)^18–20^, whilst other parts, such as returning the foraged material to a safe location are likely driven by intrinsic drives not influenced by external rewards. We therefore implemented an hybrid model where the portion of the task involving navigating to the forage location and foraging for material was learned, but the task of returning that material to the home location was controlled by intrinsic behavioural rules.

### A hybrid model successfully captures the effect of effort and an aversive environment

The results of experiments with the curriculum learning model indicate that learning every stage of the foraging sequence (navigation to forage location, act of foraging, returning foraged material to home) is not only unstable, but also results in over-learning, making it difficult to observe changes to behaviour as a result of changes in the reward function. It is often argued, however, that animals do not need to learn every phase of foraging behaviour^21,22^. We propose that, whilst finding new forage locations likely involves learning, returning foraged material is instinctive, requiring no reinforcement. In support of this conclusion, research has identified that animals do not need to learn the route back to their home location following exploration due to their ability to rely on path integration^23,24^. Given evidence that RL models explain a range of observed foraging behaviours^25–27^, as well as links between foraging behaviours and dopamine^28–30^, we assume that the behaviours of navigating to forage locations are acquired through RL mechanisms. We therefore designed a hybrid model which learned to navigate to the forage location and obtain forage material, but would then teleport the agent back to the home location to automatically drop the material, thus simulating the intrinsic drive to return foraged material to a safe location. The hybrid model was tested in a similar way as the curriculum learning model. As can be seen in Figures 2B & D, both increased effort (Kruskal Wallis: *H* = 202.435, *df* = 3, *p* =*<* 0.001, *η*^2^ = 0.841, (Effort = 50, *p <* 0.001) (Effort = 100, *p <* 0.001)) and increased aversiveness (Kruskal Wallis: *H* = 208.824, *df* = 3, *p* =*<* 0.001, *η*^2^ = 0.868, (Aversion = 1, *p* = 0.005) (Aversion = 5, *p <* 0.001) (Aversion = 10, *p <* 0.001)) reduced the number of forage bouts. Given there was no need to adjust the sharpness of the decision boundary between actions (i.e. softmax temperature), we argue that this model may be a better fit to the behaviours observed in Xeni et al. ^7^ .

### Dissociable drug mechanisms result in divergent effects on reward-seeking behaviour which is more pronounced in the EBF

In classical EfR tasks, both methylphenidate and amphetamine have been shown to increase the frequency of high-effort for high-reward behaviours^9,31,32^. However, Xeni et al. ^7^ observed that methylphenidate and amphetamine impacted learning in the EBF task in different ways. In order to capture this in our model, we hypothesised two dissociable models of drug-action. Namely, that whilst both methylphenidate and amphetamine increase reward sensitivity via a heightened learning rate^33,34^, only amphetamine also introduces noise into the TD(0) error term (*δ*)^8^.

Methylphenidate is primarily a dopamine reuptake inhibitor – it reduces the availability of dopamine transporters, thereby increasing the amount of extracellular dopamine following the natural release of dopamine^35–37^. Howlett et al. ^33^ captured this effect as an increase in the learning rate of a computational RL model. While performing a probabilistic learning task, healthy human males were administered a dose of methylphenidate. The researchers then calculated an analogue of the participants’ learning rate by measuring the change in behaviour following an error. They also fit the observed behaviours to a Rescorla-Wagner model with 2 free parameters, the learning rate and the inverse temperature of the decision function. They found that subjects who had received methylphenidate exhibited higher learning rates than those in the placebo group. A similar effect was observed by Westbrook et al. ^34^ who reported that methylphenidate increased the learning rate for human participants during a task requiring participants to learn the correct button-press response to give for each of a set of stimuli based on reward feedback. We chose to similarly model the effects of dopamine reuptake inhibition, an effect of both methylphenidate and amphetamine, as an increase in the learning rate. Amphetamine has similarly been observed to act as a dopamine reuptake inhibitor^38^. Additionally, amphetamine stimulates dopamine release, causing dopamine release to occur outside of otherwise ‘natural’ occurrences (e.g. when associated with a reward)^39^. Werlen et al. ^8^ showed that, dopaminergic signalling in the nucleus accumbens (correlating with Reward Prediction Error) were disrupted following acute doses of amphetamine, and specifically observed a reduction in the relationship between dopamine signalling and RPEs after amphetamine administration. Together these results suggest that amphetamine is associated with RPE-like signalling independent of actual rewards. Since the TD error signal resembles RPEs^40^, this effect of amphetamine can be captured by introducing noise into the TD error term.

Assuming that both the EfR and EBF tasks are subject to the same TDRL processes, there are two main differences between the tasks. The first is the involvement of additional, intrinsic processes in the EBF but not the EfR tasks. The second is the task temporal horizon. By temporal horizon we refer to the smallest number of actions or sensory states in-between an agent’s starting condition and the task goal. For example, in the simulated EfR task, the agent is required to make 4 nose-pokes to obtain a reward. Thus the temporal horizon is characterised by the 4 nose-poke actions. For the simulated EBF task, the shortest path from the agent’s start location and depositing the key in their home location is 9 actions/states. Thus the EBF task has a much longer temporal horizon than the EfR. We believe that it is this difference in temporal horizon that allows the difference in drug effects to emerge. With longer horizons, any noise or randomness injected into the error signal (i.e. via amphetamine) may result in learning incorrect associations with more states, thereby disrupting the learned behaviour. However, with short horizon tasks such as the EfR, there are fewer states about which to learn, thus restricting error propagation. Additionally, it is easier and quicker to re-visit the rewarding state and thereby provide corrective feedback. We compare both the curriculum and hybrid models to an agent trained to solve the EfR task in order to assess this hypothesis.

These results demonstrate that the curriculum model (Figure 3A) was unable to capture the effects observed by Xeni et al. ^7^ – no significant differences between conditions were observed (Kruskal Wallis: *H* = 2.413, *df* = 2, *p* = 0.299, *η*^2^ = *−*0.003). In contrast, the hybrid model (Figure 3B) successfully produced the observed effect of amphetamine whereby the presence of amphetamine led to a reduction in the average number of rewards obtained across the 60 independent trials ( Median = 28.0, IQR = (14.0 *−* 65.25)) compared to baseline (Median = 62.5, IQR = (58.0 *−* 66.25)), (*p* = 0.041). The hybrid model also captured the effect of methylphenidate whereby the animals exhibited an increase in reward-seeking behaviour (methylphenidate: Median = 77.5, IQR = (74.0 *−* 80.25), (*p <* 0.001)). Importantly, the model solving the EfR task (Figure 3C) was able to produce the increase in high-effort behaviours in response to both methylphenidate (Median = 88.0, IQR = (87.0 *−* 89.0), (*p* =*<* 0.001)), and amphetamine (Median = 80.5, IQR = (52.5 *−* 88.25), (*p* = 0.023)) compared to baseline ( Median = 72.0, IQR = (68.0 *−* 78.0)).

**Figure 3.**
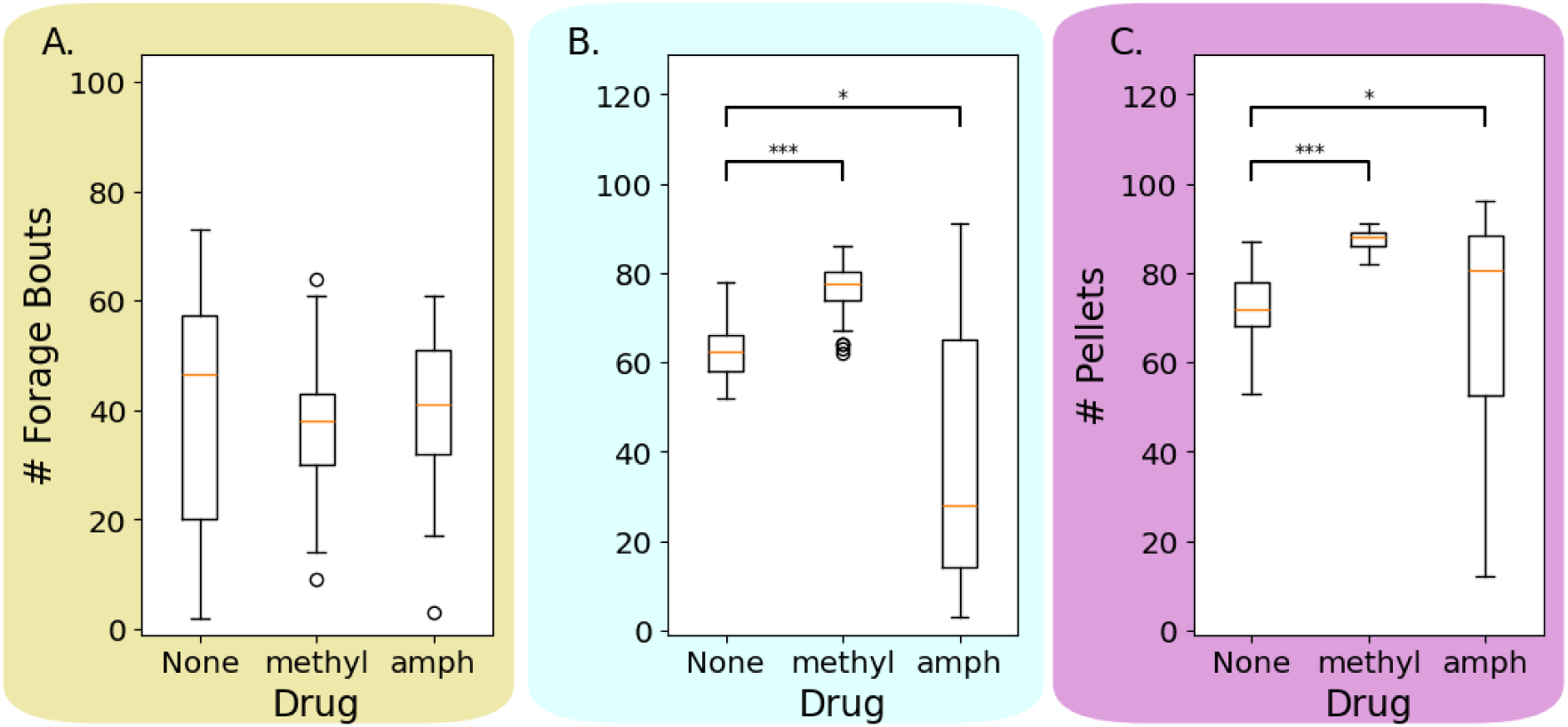
Effects of methylphenidate and amphetamine depend on task and model type. Boxplot showing the number of forage bouts or pellets obtained by each model over 60 independent trials in each drug condition. A. Curriculum model, B. Hybrid model, C. EfR task. \**p <* 0.05, \*\**p <* 0.01, \*\*\**p <* 0.001

Whilst further work is needed to fully confirm this account, we highlight that the different behavioural outcomes of the two drug mechanisms has important implications for our understanding and development of drug interventions. For example, whilst treating apathy syndrome with drugs with similar mechanisms as amphetamine may result in behavioural improvement in short-horizon motivation, the proposed models predict detriments for long-horizon motivation. In practical terms, this finding highlights the importance of testing the impact of drug interventions for apathy on a range of different tasks. Investigations into how dopaminergic drugs modulate motivation have adopted tasks with fairly immediate rewards^41,42^. Performance on these tasks is often interpreted to imply that the drug may provide potential improvements on real-world tasks, including multi-step tasks that require planning and delayed gratification (e.g. work projects, and planning social events). The present findings provide motivation for filling this evidence gap.

### TDRL-based models of drug action produce amphetamine dose-response curve predictions

We can use the computational models that successfully captured behaviours in the EfR and EBF tasks (TDRL alone and the novel hybrid model respectively) to produce testable predictions. Namely, concerning dose-response curves in each of the two tasks. To construct these curves, we sampled 40 amphetamine doses on a log scale ranging from 6.86*e −* 06 - 2. We also included dose = 0. These doses were treated as the value for the dopamine variable in the networks. We then ran the hybrid model solving the EBF task, and the simple RL model solving the EfR task for 60 independent trials at each amphetamine dose level. We collected the number of forage bouts and pellets per trial respectively and averaged the results.

The results can be seen in Figure 4. The vertical dashed line indicates a simulated dopamine level of 0.6 which was the value we chose when simulating the effects of amphetamine in the previous experiment. What this plot reveals is that, for the EfR task, once amphetamine = 0.6, the number of pellets being obtained is still above baseline (where amphetamine = 0 (i.e. the leftmost data point)). In contrast, for the hybrid model solving the EBF task, an amphetamine level of 0.6 results in fewer forage bouts than when amphetamine = 0.

**Figure 4.**
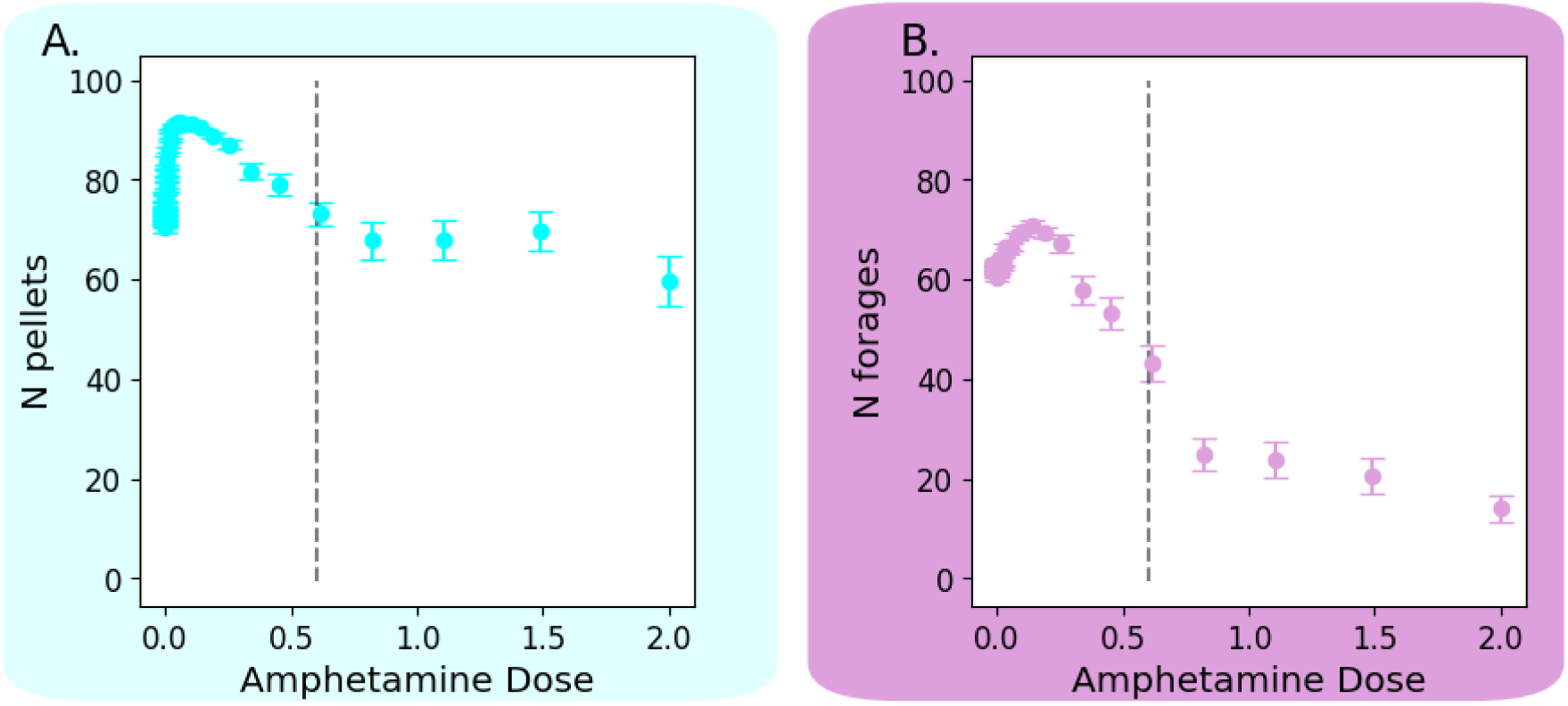
Predicted dose-response curves for increasing dopamine availability via amphetamine. Error-bar plot showing the average number of forage bouts or pellets obtained by each model over 60 independent trials in each dopamine condition. A. Hybrid modelEBF Task, B. EfR Task. Error-bars represent standard error around the mean.

What we can predict from these plots is that small doses of amphetamine should lead to an increase in reward-seeking behaviour regardless of task. As the dose increases, this facilitatory effect will eventually turn to a detrimental one, incurring a decrease in reward-seeking. The dose at which the effect changes from facilitatory to detrimental differs based on the task, with the EBF having a much smaller range of doses where any increase in reward-seeking can be observed, and a much steeper slope into detrimental effects. By mapping the simulated amphetamine doses to actual doses, future research can directly test these predictions in animal models.

## Conclusion

A primary outcome of the present work is the realisation that the TDRL mechanisms we adopted were not sufficient to produce the types of foraging behaviours seen in animals. Whilst we were able to have a model capable of learning the EBF using only TDRL (the curriculum model), this model was not able to reproduce behaviours observed in mice without significant adaptations to the strength of the learned policy. Instead we found that a hybrid model which combined learning to navigate to the forage location and forage there, with an intrinsic drive to return to the home location once forage material was obtained, was much better at accounting for the observed behaviours. This suggests that, whilst TDRL can account for parts of the foraging sequence, it is not necessarily relied on for the whole sequence. Thus, we highlight the necessity of establishing, not only the individual mechanisms driving behaviour, but also the potential interplay, and separation, between multiple mechanisms (i.e. learning and intrinsic drives).

In order to validate the developed model, we sought to provide an account for the divergence in dopaminergic drug effects observed in^7^. In EfR tasks, dopaminergic drugs such as methylphenidate and amphetamine have been shown to increase reward-seeking behaviour, indicating an important role of dopamine in motivation and thereby supporting their use as therapeutic interventions for disorders such as apathy. However, the EBF revealed that, whilst methylphenidate increases foraging behaviours, amphetamine leads to a decrease in overall quantities of material foraged^7^. In the present work, we successfully captured this observation in a computational model that incorporated, but did not depend solely on, reinforcement learning. We account for the observed divergence in drug effects as an interaction between the drug’s mechanisms of action (i.e. whether they interact with naturally occurring dopamine, or stimulate dopamine release), and the temporal horizon (i.e. the number of actions/decisions between the starting condition and the reward) of the task. Specifically, short horizon tasks such as the EfR are not sufficient for revealing the interference incurred via drug-induced dopamine release, whereas long horizon tasks such as the EBF are.

Finally, we were able to apply the developed hybrid and EfR models to produce novel, testable hypotheses regarding the dose-response curves of amphetamine on motivated behaviours. Future work should investigate these predictions in biological agents.

### Limitations

One limitation of this work lies in how we operationalise motivation, i.e. what behaviours we measure. In Xeni et al. ^7^ , the researchers measure the amount of nesting material foraged and use that as an indicator of motivation. In the simulated EBF task developed and used here, we instead measure the number of forage bouts. This is mostly due to a restriction in the simulated task design - the agent was only able to retrieve 1 key at a time. In designing the task this way we made the assumption that the mice^7^ carried the same volume of material per forage bout, but that the experimental manipulations altered the frequency of forage bouts. This assumption should be verified in future work. If it is not the case, and instead the mice carried more material per bout, the simulated task would need to be altered to allow mice to forage multiple times in a row before choosing to return to the home location. Such a redesign to increase the behavioural fidelity of the model may provide insights into the nature of the reward signal driving behaviour. For instance, the simulated task would need to incorporate a carrying capacity which would likely also involve an effort penalty such that carrying more material is more effortful. Learning the task would therefore involve striking a balance between the reward of foraging and storing material with the effort exerted in retrieval. This could be useful in highlighting individual value systems. Agents that primarily value the act of foraging, but not the foraged material itself, might spend a long time foraging but not return much of the material to the home location. At the same time, if the perceived effort of travelling between locations is greater than the effort of carrying material, agents might be expected to perform fewer forage bouts. In both instances, fewer forage bouts are observed and less material shuttled, but as the result of very different cost-benefit analyses. Extensive research identifying sources of motivation and effort and their impact on behaviour is needed to establish such a model. The current work provides a strong foundation and justification for such work.

A further limitation is that these computational models are relatively simplistic and fail to account for neurotransmitters other than dopamine, or interactions with the rest of the brain. For example, amphetamine is known to also influence serotonin and noradrenaline release and reuptake^43–45^. The neurotransmitter serotonin also plays an important role in motivation and reward-based learning^14,46^. It is therefore likely that, by not including the dynamics of the effects of amphetamine and methylphenidate on serotonin and other neurotransmitters, we have missed some nuances or obscured the true mechanisms behind the observed behaviours. A more complete model would provide further insights. However, the work done here provides valuable insights into the possible mechanisms driving the observed distinctions between drug effects, establishing testable hypotheses for evaluating an account of novel observations.

We further note that the effects of methylphenidate on learning rate reported in^33,34^ were both context-dependent. The effect reported in^33^ was true only in the first testing session, disappearing such that there was no difference between the treatment and placebo groups in the second session. The effect of methylphenidate reported in^34^ was primarily true for participants with a high dopamine synthesis capacity. For the purposes of this investigation we adopted this relatively simplistic model of methylphenidate action, as it seemed the minimal required change to capture increased reward sensitivity due to dopamine reuptake inhibition, and did not require knowledge of potentially interacting variables (e.g. dopamine synthesis capacity). It is the task of future work to characterise, in full, the computational impacts of methylphenidate.

Finally, we note that another important distinction between the EfR and EBF tasks are the sources of motivation. In the case of the EfR, behaviours are extrinsically motivated by the delivery of food/water whilst the animal is experiencing deprivation. The EBF task, on the other hand, arguably relies on intrinsic motivation, especially since the original study demonstrated that de-valuing the bedding material had no impact on the tendency to perform effortful foraging^7^. Given that RL algorithms compound all reward signals into a single value, mirroring the dopaminergic RPE signal, it is difficult to disentangle the relative impacts of different sources of motivation. Thus whilst exploring the mechanistic distinction between intrinsic and extrinsic motivation is beyond the scope of the current work, we highlight that it potentially contributes to the observed divergence in drug effects and therefore warrants further investigation.

## RESOURCE AVAILABILITY

### Lead contact

Requests for further information and resources should be directed to and will be fulfilled by the lead contact, Dr Madeleine Bartlett.

### Materials availability

New materials generated by this study include the simulated Effort for Reward and Effort Based Foraging tasks. Code for these tasks will be made available on GitHub following journal submission.

### Data and code availability

Data and code will be made available after journal submission.

## ACKNOWLEDGMENTS

This work was funded by National Research Council of Canada’s Artificial Intelligence for Design program via grant AI4D-151-1. M.J was funded by the SWBio BBSRC Doctoral career development fund. The authors thank all members of their labs for their support.

## AUTHOR CONTRIBUTIONS

Conceptualization, M.B. and M.J.; methodology, M.B. and M.J.; investigation, M.B. and M.J.; writing – original draft, M.B. and M.J.; writing – review & editing, P.M.F., T.C., J.O., M.B. and M.J.; funding acquisition, T.C. and J.O.; resources, T.C. and J.O.; supervision, M.J., P.M.F., T.C. and J.O..

## STAR METHODS

### Key resources table

| REAGENT or RESOURCE | SOURCE | IDENTIFIER |
| --- | --- | --- |
| **Deposited data** |  |  |
| Analysed data | This paper | N/A |
| **Software and algorithms** |  |  |
| Custom MiniGrid environment | GitHub | Custom Effort for Reward tas |
| GitHub | Reinforcement learning models | GitHub |
| Python scripts for running experiments | GitHub | Python scripts for data analy |
| GitHub |  |  |

### Method details

#### Reinforcement Learning Model

We implemented a simple policy-gradient actor-critic network using look-up tables to store the learned value function and policy. For solving the EBF task we store the value function as a 7 *×* 7 *×* 4 array *V* , using the one-hot encoding of the state *S_t_* (also a 7 *×* 7 *×* 4 array) to index the value function and retrieve or update the stored value. Similarly, *z* stores the 7 *×* 7 *×* 4 *×* 5 vectorised policy. We index the policy using the one-hot encoded action *A_t_*, retrieving or updating the 5-element vector containing the values or logits of the actions given our current state *S_t_*. For the EfR task, the value function was stored in an *N* vector, and the policy in an *N ×* 2 array.

In all cases, we employed the temporal difference TD(0) learning rule^16^ for updating the values stored in the look-up tables. Under this algorithm, at each time step of the environment we calculate the one-step return as:

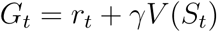

where *r_t_* is the reward obtained by moving into state *S_t_*, *V* [*S_t_*] is the value associated with that state, and *γ* is a discount term. This is then used in calculating the TD(0) error term *δ*,

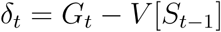

where *V* [*S_t−_*_1_] is the value associated with the previous state the agent was in. We then update the value of the previous state (*V* [*S_t−_*_1_]) stored in the look-up table according to

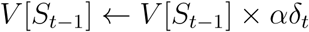

where *α* is the learning rate. The policy is similarly stored in a look-up table. For each state *S*, we store the learned logits *z*[*S*] for each of the 5 available actions. We also update the policy by updating the logits for the actions stored in the policy look-up table,

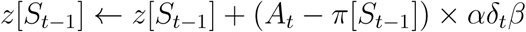

where *A_t_* is a one-hot vector encoding the action that was selected (i.e. *π_i_*(*S_t−_*_1_) is the probability of action *i* at state *S_t−_*_1_ according to a softmax policy), and *β* is a discount term.

Values for the hyperparameters *α*, *γ*, and *β* were identified through manual optimisation and are shown in Table 1. Different values were selected depending on which reward function was used to train the model (curriculum learning vs. hybrid learning & intrinsic). We describe these reward functions in *Training Procedures*.

**Table 1:**
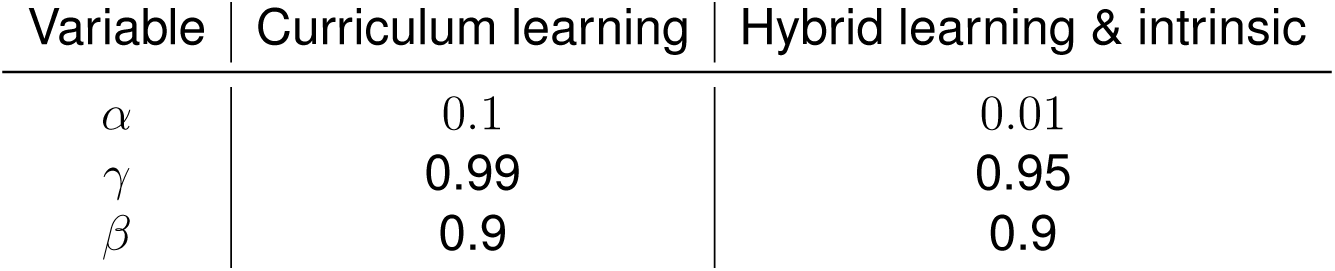
Learning rule parameter values used to train network under each reward function condition. Values were identified via a manual optimisation search.

| Variable | Curriculum learning | Hybrid learning & intrinsic |
| --- | --- | --- |
| $\alpha$ | 0.1 | 0.01 |
| $\gamma$ | 0.99 | 0.95 |
| $\beta$ | 0.9 | 0.9 |

#### Effort for Reward Task

In the EfR task, the agent is presented with two action options: (1) a low-effort action whereby the agent incurs no cost and immediately obtains a reward of 0.1, (2) a high-effort action, incurring a cost of *−*0.2 to the total reward obtained per time step. After *N* repetitions of the high-effort action, the agent receives a reward of 1.0. This task replicates the EfR task used in Xeni et al. ^14^ , where animals are placed in an environment where they can perform an effortful action to obtain sugar pellets (high value reward), but also have access to freely available chow (low value reward). The agent can be in one of *N* states whereby *N* is the number of high-effort actions required to obtain the high-value reward.

#### Effort Based Foraging Task

For the simulated version of the EBF task, we developed a custom environment using the Farama Foundation MiniGrid environments^17^ as a base. The bespoke environment is depicted in Figure 1. We adapted a simple, empty 7 *×* 7 grid world to contain a key object, representing the forage material, in the bottom right-hand corner. The agent’s ‘home location’ was situated in the top left-hand corner. In each time step the agent would choose to take one of 5 actions: *turn left*, *turn right*, *move forward*, *pick up*, or *drop*. When the agent obtained a key, a new key was immediately placed in the same location, allowing the agent to continuously forage without having to reset the environment.

We felt that it was important to allow the agent to behave in a continuous manner, without having the environment reset. This was to establish that the agent had learned the task properly - learning to navigate to the forage location, obtain a key, return it to home and then navigate back to the forage location to obtain another key. The agent can be in one of 7 *×* 7 *×* 4 *×* 2 states corresponding to the agent’s x location, y location, orientation, and whether or not it is holding a key, respectively.

For both tasks, the state space was represented using a one-hot vector, and the value function and policy were stored in look-up tables.

#### Training Procedures

Three different training procedures were used. Once the agent was trained to successfully solve the task, the learned value function and policy were saved and stored to be used when testing the agents under different conditions.

#### EfR Training - Baseline RL

We trained a pure RL agent on the EfR task using the learning algorithm and reward function described above. To mirror the training procedure in^14^, for the first 30 learning trials we trained the agent to perform 2 high-effort actions for the high-value reward. We then increased this to 4 high-effort actions for the remaining 30 learning trials. To ensure that the agent had properly learned the task, the learned policy was manually reviewed to determine that the agent had appropriately learned to take the high-effort action in each state.

#### EBF Training - Curriculum RL

In order to train a pure RL agent on the EBF task we implemented curriculum learning, breaking the task up into different stages. For the first 150 learning trials, the agent was set to always be holding forage material. The agent started at a location facing the forage location, and was trained to return to the home location and use the *drop* action to drop the key in the home square and obtain a reward. For the next 100 trials, the agent started in the same location but was no longer initialised as holding forage material. The agent was rewarded for using the *pick up* action to obtain the forage material at the forage location, and then again for depositing the key in the home square. Finally, for learning trials 250-600, the agent was initialised in the home location and trained to navigate to the forage box, obtain the forage material, then return it to the home box. We also implemented a ‘continuous’ phase where the environment was no longer reset between forage trials. Instead, the agent was given a total of 100 time steps in which to behave in the environment.

Thus, the agent is exposed to a total of 3 separate reward signals. The first is the reward for dropping forage material in the home box, *R_h_* = 0.8. The second is the reward for performing the *pick up* action in order to forage, *R_f_* = 2.0. The final reward signal is received in tandem with the forage reward and is a signal indicating the effort required to obtain the forage material, *R_e_* = *−*0.5.

It should be noted that correctly learning this task was not robust to randomness in the initialisation of the network. In many cases the agent would learn to retrieve the forage material but, once it returned it to the home location, it would stay in its same location and repeatedly pick-up and drop the same object. This did not provide any additional reward signals and so was not a reinforced behaviour. However, because of this it was necessary to ensure that the agent had learned the appropriate policy function before saving that function for further experiments. We did this by checking whether the agent achieved an epsiodic reward of *>* 200 by the end of the learning phase.

#### EBF Training - Hybrid Learning & Intrinsic

The results of experiments with the curriculum learning model indicated that learning the whole task from end-to-end was not only unstable, but also resulted in over-learning, making it difficult to observe changes to behaviour as a result of changes in the reward function. Whilst it is arguably necessary for animals to learn how to find new forage locations, and to learn when that location’s resources are depleted, it is arguably not necessary for animals to learn to return forage material to their home location. This may instead be an intrinsically driven behaviour requiring no reinforcement through reward signals. We therefore designed a hybrid model which learned the first portion of the task – to navigate to the forage location and obtain forage material – but which would then ‘teleport’ the agent back to the home location where it automatically dropped the material. This teleportation was used as an analog for animals’ reliance on path integration, and not learning, to return to their original locations^23,24^. We did not require curriculum training to train this model, the agent was instead given 900 identical learning trials. The agent would start in the home location and had to learn to navigate to the forage location where it would earn a reward of 1.0 for obtaining forage material, with a negative reward of *−*0.5 for exerting energy to take the ’pick-up’ action. In order to simulate the intrinsic drive to return forage material to a nest the agent would then be teleported back to the home location. We prevented overfitting by employing early stopping procedures.

## 1 Testing Conditions

In all experiments each condition was tested over 60 independent runs and results were averaged.

### Effort

For the first experiment we wanted to see whether increasing the required amount of effort would alter the agent’s behaviour in the EBF task. We took the trained agent and subjected it to 4 different effort conditions. We operationalised increased effort as an increase in the amount of negative reward associated with the foraging action. During training, the negative reward for effort was *−*0.5. In the 4 effort conditions, we increased effort in the EBF task by multiplying this negative feedback by *×*1, *×*10, *×*50, and *×*100.

### Aversion

We also wanted to examine the impact of increasingly aversive environments on agent behaviour. This was as a proxy for making the environment larger, as in Xeni et al. ^7^ . At every time step where the agent was away from the home location, the agent received a negative reward which we refer to as the aversion score. We again created 4 conditions, testing aversion scores of 0, 1, 5, and 10.

### Acute Pharmacology

We further tested the hypothesis that the disparity in drug effects observed by Xeni et al. ^7^ was due to a difference in tonic vs. phasic drug interactions. Namely, given that methylphenidate is a dopamine reuptake inhibitor, we propose that this drug only effects naturally released dopamine. However, amphetamine is both a reuptake inhibitor and a dopamine stimulant. We therefore propose that, as well as influencing reactivity to naturally released dopamine, amphetamine also incurs tonic effects via the release of dopamine regardless of the presence of RPE eliciting stimuli. In particular, we propose that amphetamine has the effect of diluting ‘true’ RPEs. We model these effects in the following ways.

Based on previous work suggesting that methylphenidate increases sensitivity to rewards and reinforcing stimuli, we chose to model the effect of this drug as an increase in the learning rate^33,34,47^. Increased DA levels are associated with heightened sensitivity to rewards. By elevating the learning rate, we capture this sensitivity by forcing the agent to learn more quickly. Amphetamine has similarly been shown to increase reward sensitivity^9,48–50^ so we also adopt this approach to modelling its influence on phasic dopamine signals. However, in regards to amphetamine’s tonic influence obscuring learning signals, research has indicated that amphetamine can disrupt reward prediction error signals. For example Werlen et al. ^8^ demonstrated that, following amphetamine administration, it is no longer possible to distinguish between the average

RPE in response to a low-value vs. a high-value reward cue. We therefore additionally model amphetamine by introducing a background randomness term *ɛ* into the TD error equation:

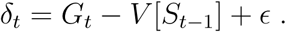

### Quantification and statistical analysis

To calculate the number of completed forage bouts, we counted the number of times the agent received a non-zero reward indicating that they had either dropped the forage material at the home location (curriculum model), or picked up forage material at the forage location (hybrid model). We similarly calculated the number of pellets obtained by counting every 4th nose-poke. We first tested the data for normality using the Shapiro-Wilk test. Comparison’s between condition were made using the Kruskal-Wallis non-parametric test. We collected results from 60 independent trials in each condition for these analyses and report the medians and interquartlie ranges of the results. The significance of these tests is indicate in Figures 2 and 3. Significance was defined as a p-value *>* 0.5.

To generate the predicted dose-response curves shown in Figure 4, we collected the results from 60 independent trials in each amphetamine dose condition. We then plotted the mean and standard error around the mean in an error-bar plot.

## References

1. Grima, L.L., Haberkern, H., Mohanta, R., Morimoto, M.M., Rajagopalan, A.E., and Scholey, E.V. (2025). Foraging as an ethological framework for neuroscience. Trends in Neurosciences.

2. Chase, T.N. (2011). Apathy in neuropsychiatric disease: diagnosis, pathophysiology, and treatment. Neurotoxicity research 19, 266–278.

3. Benoit, M., Andrieu, S., Lechowski, L., Gillette-Guyonnet, S., Robert, P.H., and Vellas, B. (2008). Apathy and depression in alzheimer’s disease are associated with functional deficit and psychotropic prescription. International Journal of Geriatric Psychiatry: A journal of the psychiatry of late life and allied sciences 23, 409–414.

4. Den Brok, M.G., van Dalen, J.W., van Gool, W.A., Moll van Charante, E.P., de Bie, R.M., and Richard, E. (2015). Apathy in parkinson’s disease: a systematic review and meta-analysis. Movement Disorders 30, 759–769.

5. Salamone, J.D., Steinpreis, R., McCullough, L., Smith, P., Grebel, D., and Mahan, K. (1991). Haloperidol and nucleus accumbens dopamine depletion suppress lever pressing for food but increase free food consumption in a novel food choice procedure. Psychopharmacology 104, 515–521.

6. Richardson, N.R., and Roberts, D.C. (1996). Progressive ratio schedules in drug self-administration studies in rats: a method to evaluate reinforcing efficacy. Journal of neuroscience methods 66, 1–11.

7. Xeni, F., Marangoni, C., and Jackson, M.G. (2024). Validation of a non-food or water motivated effort-based foraging task as a measure of motivational state in male mice. Neuropsychopharmacology 49, 1883–1891.

8. Werlen, E., Shin, S.L., Gastambide, F., Francois, J., Tricklebank, M.D., Marston, H.M., Huxter, J.R., Gilmour, G., and Walton, M.E. (2020). Amphetamine disrupts haemodynamic correlates of prediction errors in nucleus accumbens and orbitofrontal cortex. Neuropsychopharmacology 45, 793–803.

9. Marangoni, C., Tam, M., Robinson, E.S., and Jackson, M.G. (2023). Pharmacological characterisation of the effort for reward task as a measure of motivation for reward in male mice. Psychopharmacology 240, 2271–2284.

10. Drui, G., Carnicella, S., Carcenac, C., Favier, M., Bertrand, A., Boulet, S., and Savasta, M. (2014). Loss of dopaminergic nigrostriatal neurons accounts for the motivational and affective deficits in parkinson’s disease. Molecular psychiatry 19, 358–367.

11. Magnard, R., Vachez, Y., Carcenac, C., Krack, P., David, O., Savasta, M., Boulet, S., and Carnicella, S. (2016). What can rodent models tell us about apathy and associated neuropsychiatric symptoms in parkinson’s disease? Translational psychiatry 6, e753–e753.

12. Mitchell, R.A., Herrmann, N., and Lanctôt, K.L. (2011). The role of dopamine in symptoms and treatment of apathy in alzheimer’s disease. CNS neuroscience & therapeutics 17, 411– 427.

13. Sgambato-Faure, V., and Tremblay, L. (2018). Dopamine and serotonin modulation of motor and non-motor functions of the non-human primate striato-pallidal circuits in normal and pathological states. Journal of Neural Transmission 125, 485–500.

14. Xeni, F., Marangoni, C., Lin, L., Robinson, E.S., and Jackson, M.G. (2025). Conditioned versus innate effort-based tasks reveal divergence in antidepressant effect on motivational state in male mice. Neuropsychopharmacology pp. 1–11.

15. Schultz, W., Dayan, P., and Montague, P.R. (1997). A neural substrate of prediction and reward. Science 275, 1593–1599.

16. Sutton, R.S., and Barto, A.G. (2018). Reinforcement learning: An introduction. MIT press.

17. Chevalier-Boisvert, M., Dai, B., Towers, M., de Lazcano, R., Willems, L., Lahlou, S., Pal, S., Castro, P.S., and Terry, J. (2023). Minigrid & miniworld: Modular & customizable reinforcement learning environments for goal-oriented tasks. CoRR abs/2306.13831.

18. Jiang, W.C., Xu, S., and Dudman, J.T. (2022). Hippocampal representations of foraging trajectories depend upon spatial context. Nature neuroscience 25, 1693–1705.

19. Goldshtein, A., Handel, M., Eitan, O., Bonstein, A., Shaler, T., Collet, S., Greif, S., Medellín, R.A., Emek, Y., Korman, A., et al. (2020). Reinforcement learning enables resource partitioning in foraging bats. Current Biology 30, 4096–4102.

20. Dubois, T., Pasquaretta, C., Barron, A.B., Gautrais, J., and Lihoreau, M. (2021). A model of resource partitioning between foraging bees based on learning. PLOS Computational Biology 17, e1009260.

21. Hall-McMaster, S., and Luyckx, F. (2019). Revisiting foraging approaches in neuroscience. Cognitive, Affective, & Behavioral Neuroscience 19, 225–230.

22. Frankenhuis, W.E., Panchanathan, K., and Barto, A.G. (2019). Enriching behavioral ecology with reinforcement learning methods. Behavioural Processes 161, 94–100.

23. Whishaw, I.Q., Hines, D.J., and Wallace, D.G. (2001). Dead reckoning (path integration) requires the hippocampal formation: evidence from spontaneous exploration and spatial learning tasks in light (allothetic) and dark (idiothetic) tests. Behavioural brain research 127, 49–69.

24. Etienne, A.S., and Jeffery, K.J. (2004). Path integration in mammals. Hippocampus 14, 180–192.

25. Montague, P.R., Dayan, P., and Sejnowski, T.J. (1996). A framework for mesencephalic dopamine systems based on predictive hebbian learning. Journal of neuroscience 16, 1936–1947.

26. Wispinski, N.J., Butcher, A., Mathewson, K.W., Chapman, C.S., Botvinick, M.M., and Pilarski, P.M. (2022). Adaptive patch foraging in deep reinforcement learning agents. arXiv preprint arXiv:2210.08085.

27. Niv, Y., Joel, D., Meilijson, I., and Ruppin, E. (2002). Evolution of reinforcement learning in uncertain environments: A simple explanation for complex foraging behaviors.

28. Rutledge, R.B., Lazzaro, S.C., Lau, B., Myers, C.E., Gluck, M.A., and Glimcher, P.W. (2009). Dopaminergic drugs modulate learning rates and perseveration in parkinson’s patients in a dynamic foraging task. Journal of Neuroscience 29, 15104–15114.

29. Le Heron, C., Kolling, N., Plant, O., Kienast, A., Janska, R., Ang, Y.S., Fallon, S., Husain, M., and Apps, M.A. (2020). Dopamine modulates dynamic decision-making during foraging. Journal of Neuroscience 40, 5273–5282.

30. Howe, M.W., Tierney, P.L., Sandberg, S.G., Phillips, P.E., and Graybiel, A.M. (2013). Prolonged dopamine signalling in striatum signals proximity and value of distant rewards. nature 500, 575–579.

31. Floresco, S.B., Tse, M.T., and Ghods-Sharifi, S. (2008). Dopaminergic and glutamatergic regulation of effort-and delay-based decision making. Neuropsychopharmacology 33, 1966–1979.

32. Ecevitoglu, A., Rotolo, R.A., Edelstein, G.A., Goldhamer, A., Mitola, M., Presby, R.E., Yu, A., Pietrorazio, D., Zorda, E., Correa, M. et al. (2025). Effort-related motivational effects of methylphenidate: Reversal of the low-effort bias induced by tetrabenazine and enhancement of progressive ratio responding in male and female rats. Neuropharmacology 269, 110345.

33. Howlett, J.R., Huang, H., Hysek, C.M., and Paulus, M.P. (2017). The effect of single-dose methylphenidate on the rate of error-driven learning in healthy males: a randomized controlled trial. Psychopharmacology 234, 3353–3360.

34. Westbrook, A., van den Bosch, R., Hofmans, L., Papadopetraki, D., Määttä, J.I., Collins, A.G., Frank, M.J., and Cools, R. (2025). Striatal dopamine can enhance both fast working memory, and slow reinforcement learning, while reducing implicit effort cost sensitivity. Nature Communications 16, 6320.

35. Volkow, N.D., Wang, G.J., Fowler, J.S., Logan, J., Gerasimov, M., Maynard, L., Ding, Y.S., Gatley, S.J., Gifford, A., and Franceschi, D. (2001). Therapeutic doses of oral methylphenidate significantly increase extracellular dopamine in the human brain. The Journal of neuroscience 21, RC121.

36. van den Bosch, R., Lambregts, B., Määttä, J., Hofmans, L., Papadopetraki, D., West-brook, A., Verkes, R.J., Booij, J., and Cools, R. (2022). Striatal dopamine dissociates methylphenidate effects on value-based versus surprise-based reversal learning. Nature Communications 13, 4962.

37. Lan, D.C., and Browning, M. (2022). What can reinforcement learning models of dopamine and serotonin tell us about the action of antidepressants? Computational Psychiatry 6, 166.

38. Jones, S.R., Joseph, J.D., Barak, L.S., Caron, M.G., and Wightman, R.M. (1999). Dopamine neuronal transport kinetics and effects of amphetamine. Journal of neurochemistry 73, 2406–2414.

39. Daberkow, D., Brown, H., Bunner, K., Kraniotis, S., Doellman, M., Ragozzino, M., Garris, P.A., and Roitman, M.F. (2013). Amphetamine paradoxically augments exocytotic dopamine release and phasic dopamine signals. Journal of Neuroscience 33, 452–463.

40. Schultz, W. (2016). Dopamine reward prediction error coding. Dialogues in clinical neuroscience 18, 23–32.

41. Chong, T.T.J., Fortunato, E., and Bellgrove, M.A. (2023). Amphetamines improve the motivation to invest effort in attention-deficit/hyperactivity disorder. Journal of Neuroscience 43, 6898–6908.

42. Bogdanov, M., LoParco, S., Otto, A.R., and Sharp, M. (2022). Dopaminergic medication increases motivation to exert cognitive control by reducing subjective effort costs in parkinson’s patients. Neurobiology of Learning and Memory 193, 107652.

43. Martin, D., and Le, J.K. (2020). Amphetamine.

44. Sloviter, R.S., Drust, E.G., and Connor, J.D. (1978). Evidence that serotonin mediates some behavioral effects of amphetamine. The Journal of Pharmacology and Experimental Therapeutics 206, 348–352.

45. Sulzer, D., Sonders, M.S., Poulsen, N.W., and Galli, A. (2005). Mechanisms of neurotransmitter release by amphetamines: A review. Progress in Neurobiology 75, 406–433. URL: https://www.sciencedirect.com/science/article/pii/S0301008205000432. doi: 10.1016/j.pneurobio.2005.04.003.

46. Izquierdo, A., Carlos, K., Ostrander, S., Rodriguez, D., McCall-Craddolph, A., Yagnik, G., and Zhou, F. (2012). Impaired reward learning and intact motivation after serotonin depletion in rats. Behavioural brain research 233, 494–499.

47. Rostami Kandroodi, M., Cook, J.L., Swart, J.C., Froböse, M.I., Geurts, D.E., Vahabie, A.H., Nili Ahmadabadi, M., Cools, R., and den Ouden, H.E. (2021). Effects of methylphenidate on reinforcement learning depend on working memory capacity. Psychopharmacology 238, 3569–3584.

48. Soder, H.E., Cooper, J.A., Lopez-Gamundi, P., Hoots, J.K., Nunez, C., Lawlor, V.M., Lane, S.D., Treadway, M.T., and Wardle, M.C. (2021). Dose-response effects of d-amphetamine on effort-based decision-making and reinforcement learning. Neuropsychopharmacology 46, 1078–1085.

49. Wyvell, C.L., and Berridge, K.C. (2000). Intra-accumbens amphetamine increases the conditioned incentive salience of sucrose reward: enhancement of reward “wanting” without enhanced “liking” or response reinforcement. Journal of Neuroscience 20, 8122–8130.

50. Krivanek, J.A., and McGaugh, J.L. (1969). Facilitating effects of pre-and posttrial amphetamine administration on discrimination learning in mice. Agents and Actions 1, 36–42.

